# Just how big is intraspecific trait variation in basidiomycete wood fungal fruit bodies?

**DOI:** 10.1101/593517

**Authors:** Samantha K. Dawson, Mari Jönsson

## Abstract

As the use of functional trait approaches is growing in fungal ecology, there is a corresponding need to understand trait variation. Much of trait theory and statistical techniques are built on the assumption that interspecific variation is larger than intraspecific variation. This allows the use of mean trait values for species, which the vast majority of trait studies adopt. We examined the size of intra- vs. inter-specific variation in two wood fungal fruit body traits: size and density. Both coefficients of variation (CV) and Trait Probability Density analyses were used to quantify trait variation. We found that intraspecific variation in fruit body density was more than twice as variable as interspecific variation, and fruit body size was hugely variable (CVs averaged 190%), although interspecific variation was larger. Further, there was a very high degree of overlap in the trait space of species, indicating that there may be little niche partitioning at the species level. These findings show that intraspecific variation is highly important and should be accounted for when using trait approaches to understand fungal ecology. More data on variation of other fungal traits is also desperately needed to ascertain whether the high level of variation found here is typical for fungi. While the need to measure individuals does reduce the ability to generalise at the species level, it does not negate the usefulness of fungal trait measurements. There are two reasons for this: first, the ecology of most fungal species remains poorly known and trait measurements address this gap; and secondly, if trait overlap between species more generally is as much as we found here, then individual measurements may be more helpful than species identity for untangling fungal community dynamics.

## Introduction

There is a growing number of studies advocating the use of functional trait approaches to understand fungal species and communities, especially accounting for within-species variation which has been largely ignored (Behm and Kiers, 2014; Crowther *et al.*, 2014; Aguilar-Trigueros *et al.*, 2015; Dawson *et al.*, 2018). By focusing on characteristics, rather than species identities, trait methods allow a deeper understanding of how species and communities perform, function and interact with each other, other organisms, and the environment. Functional traits are increasingly applied in fungal research, revealing important insights into ecological roles or functions in communities (Bässler *et al.*, 2014; Ottosson *et al.*, 2015; Halbwachs, Simmel and Bässler, 2016) and environmental dependencies (Nordén *et al.*, 2013; Abrego *et al.*, 2017). While there has been a growing number of studies examining intraspecific variation in mycorrhizal fungi (Hazard, Kruitbos, Davidson, Mbow, *et al*., 2017; Hazard, Kruitbos, Davidson, Taylor, & Johnson, 2017; Johnson *et al.*, 2012), they largely use isolates sourced from different populations rather than co-existing individuals in the field (Hazard and Johnson, 2018). For other fungal types, particularly wood fungi, studies overwhelmingly use a mean trait value to represent an entire species, and the vast majority rely on identification books to source these values (Dawson *et al.*, 2018). There has not yet been a quantitative evaluation of intra- to inter-specific variation in these organisms, especially not based on empirical field data.

It is important in trait studies to have some idea of how intra- and inter-specific variation compares, due to the assumptions of trait theory and trait statistics. Both theory and statistics of functional traits were developed using plants as the model organism, where generally interspecific variation is larger than intraspecific variation (McGill *et al.*, 2006; Violle *et al.*, 2012). This led to the rise of using species means when undertaking analyses, which are designed assuming intraspecific variation is smaller (Messier *et al.*, 2010; Violle *et al.*, 2012; Shipley *et al.*, 2016). Mean trait statistics have practical advantages in that they allow for fewer trait measurements per species, borrowing of trait values from other studies, and make it easier to glean insights into the dynamics of large communities (Shipley *et al.*, 2016). However, even in plants this approach has been called into question both empirically and theoretically (Jung *et al.*, 2010; Messier *et al.*, 2010; Laughlin *et al.*, 2012; Shipley *et al.*, 2016). If intraspecific variation is large and mean traits for species are used, then this can lead to missing or misunderstanding the ecological mechanisms underpinning trait patterns, inappropriate null models, biases of coexistence estimation and underestimation of species survivability in a given environment (Violle *et al.*, 2012). These biases may occur even when intraspecific variation is smaller than interspecific variation, but large enough to influence trait-environment interactions (Jung *et al.*, 2010; Messier *et al.*, 2010). However measuring intraspecific variation is not only important for understanding how it compares to interspecific variation; these measurements can deepen our understanding of a species niche, response to the environment, specialisation and general ecology.

Wood-decay basidiomycete fungi are vital for ecosystem processes in forests, highly biodiverse (Heilmann-Clausen *et al.*, 2015) and no exception when it comes to lack of knowledge on fungal intraspecific variation. These fungi are the primary decomposers of wood, which in turn affects processes such as soil formation and nutrient cycles (Lonsdale *et al.*, 2007). They also support a range of other organisms, including vertebrates, invertebrates, plants and other fungi, that use wood-fungi as a source of nutrients or nesting habitat for invertebrates (Jonsell *et al.*, 1999). Despite their importance, wood-decay fungi in many countries are threatened by forestry practices (Valentín *et al.*, 2014). Therefore, gaining a deeper understanding of these species is necessary for successful management and, for some species, their survival. For example, forest management has negative effects on wood fungi with large fruit bodies that fruit late in the decomposition (Abrego *et al.*, 2017). Organism traits capturing biomass or size are fundamental in trait-environment research (McGill *et al.*, 2006). In terms of fungi, large fruit bodies have a higher chance to resist stress and survive for longer periods (Bässler *et al.*, 2014; Halbwachsv2016). Having larger, more structured and denser fruit bodies, likely also requires more resources for their production, which vary across their environments (Aguilar-Trigueros *et al.*, 2015; Bässler *et al.*, 2015; Abrego *et al.*, 2017). Fundamentally, fungal fruiting biomass also represents the investment in reproduction and dispersal, where species with larger or more structured (denser) fruit bodies generally sporulate longer and possess larger spore-producing tissue (hymenium), thus leading to higher reproductive and dispersal capacity (Bässler *et al.*, 2014, 2015; Aguilar-Trigueros *et al.*, 2015; Halbwachs *et al.*, 2016; Abrego *et al.*, 2017; Dawson *et al.*, 2018). Despite this crucial importance of fruit body size and density, studies do often not quantify these traits.

This paper examines the intra- and interspecific variation of fruit body size and density of wood-decay basidiomycetes occupying *Picea abies* logs in Northern Sweden. As an exploratory analysis, the overarching aim of this study is to build some baseline data about trait variation in wood-decay fungal communities. We were particularly interested in (a) establishing if interspecific variation is larger than intraspecific variation for these traits and (b) to quantify the variability of these fungal traits across species. Fruit body traits were chosen as they are becoming commonly used in saprotrophic fungi trait analyses (e.g. Nordén *et al.*, 2013; Abrego *et al.*, 2017). Additionally, as discussed in the previous paragraph, these traits in particular are important for fungal reproduction and dispersal. This is the first study we are aware of to quantify community intra- to inter-specific variation from field data in not just wood-decay fungi, but fungi more generally.

## Methods

### Study site

This study was conducted in isolated forest patches in Norrbotten County, within the Boreal Zone of Northern Sweden. Mean annual precipitation is ca. 400-700 mm; mean monthly temperatures range from −9 to −16°C (January) to 11-15°C (July). Sites were based on previous work examining the differences between set-aside forest patches of high biodiversity value (i.e. key habitats) surrounded by managed forests and natural forest patches surrounded by mires (Berglund and Jonsson, 2005). Patch sizes ranged from 0.1 to 12 ha and we surveyed 32 set-aside patches and 11 natural patches. Previous surveys of these sites have demonstrated that a large range of wood-decaying basidiomycete fungi can be found and there was no predicted extinction debt in the set-aside patches (Berglund and Jonsson, 2005). Patches were of moist to mesic ground condition (Hägglund and Lundmark, 1987) and moraine soil, with Norway spruce (*Picea abies*) dominating the tree layer and mainly bilberry (*Vaccinium myrtillus*) dominating the field layer.

### Survey methods

Following the methods of Berglund & Jonsson (2003; 2005), we returned to the central point of the patch and a 20 m diameter circular plot was placed around this point. Trait measurements were taken from fruit bodies occurring on the spruce log that was closest to the central point of the plot that fulfilled the following requirements: of mid decay class three or four (wood hard or starting to soften and < 50% bark remaining; McCullough 1948; Söderström 1988), at least 1 m in length and at least 10 cm diameter at the base (diameter range = 10-55 cm). If no log of decay classes three or four were present in the plot, a decay class of two (wood hard, 50% of bark remaining) fitting the other requirements was used. In the two cases where no suitable log was present inside the plot, we took the closest log outside the plot that fit the criteria. We focused on mid decay classes since they are known to host the greatest number and diversity of fruiting wood-decay basidiomycetes (Jönsson *et al.*, 2008; Nordén *et al.*, 2013).

Each Norway spruce log was first surveyed (non-destructively) to identify species and fruiting extent. We surveyed all polyporous and six corticoid wood-decaying fungi: *Asterodon ferruginosus*, *Cystostereum murrai*, *Laurilia sulcata*, *Phlebia centrifuga*, *Stereum sanguinolentum*, and *Veluticeps abietina*. The wood fungi we surveyed are important wood decomposers and are mainly confined to conifer forests. We then took measurements for three randomly selected fruit bodies per species, or all fruit bodies if there were three or less. It should be noted that we treat individual fruit bodies taken from the same log as just that: individual fruit bodies, rather than as being sourced from one or several individual fungi. The analyses to establish if they came from individual fungi were outside the scope of this study and potentially unnecessary. Although taking multiple measurements from the same log may mean that the same fungal individual was measured, this should not have a large effect on our interpretation for two reasons: first, fruit bodies are a part, or organ of the larger individual, and it is acceptable, in the trait literature, to sample organs of an individual multiple times, as long as many individuals are sampled (e.g. multiple leaves from the same tree for plants; Pérez-Harguindeguy *et al.*, 2013); and secondly, even if the same individual was sampled multiple times, this would bias our data towards underestimation of intraspecific variation, which means that the true intraspecific variation may be higher in some cases than what is presented here.

We measured two fruit body traits: fruit body size and dry fruit body density, following the methods of Dawson *et al*. (2018). For fruit body size, the maximum fruit body width, length and depth was measured with callipers (mm) and then volume was calculated assuming an elliptical shape (Dawson *et al*. 2018, but measured in cm^3^). In cases where very large resupinate fruit bodies (i.e. fruiting extending over 1 m long), or hundreds of smaller resupinate fruit bodies occurred in between larger fruit bodies (>30 cm long), we took four measurements to jointly characterise fruiting over the whole log: fruiting length was measured at equal intervals along this length (i.e. 25%, 50% and 75%) the width and depth of fruiting was measured. Cores for fruit body density were also taken at these points. For these very large resupinate fruit bodies the average of the three measurements of width and depth and the total length were used in volume calculations. For fruit body density, in the majority of cases a 12 mm push corer was used to sample the fruit body (directly through the hymenium), and the length of this core was measured at the same time to calculate sample volume (Dawson *et al*. 2018). For all thin (<1 mm thick) fruit body samples a standard of 0.5 mm was set, except in the rare case of an extremely thin sample where four cores were taken to increase the thickness to 0.5 mm. Where the hymenium of fruit bodies was smaller than the push core aperture, the entire fruit body was taken and volume measured as above. Sample volumes were then combined with the dry weights to obtain values for fruit body density (Dawson *et al*. 2018, mg/mm^3^). In the cases of large resupinate fruit bodies, the density was averaged across the three samples.

### Data analysis

Across the 43 sites and logs, a total of 239 fruit body measurements were taken. Of these, 36 were multiple measurements from single, large fruit bodies, resulting in 167 measurements of individual fruit bodies. We collected data on 22 species, however we removed species that had less than four measurements as three or fewer fruit bodies generally meant that the data was all collected from one log and we did not consider this to be representative. This removed 10 species from the analysis and brought the number of measurements down to 146. The remaining 12 species all had at least five measurements; the average number of measurements was 12 and 25 was the highest, made on the common resupinate polypore *Antrodia serialis*. Although the number of measurements per species are relatively low, five is the minimum number of replicate measurements to characterise most traits of plants (Pérez-Harguindeguy *et al.*, 2013) and it is unknown how many replicates are needed for fungi (Dawson *et al.*, 2018). For the number of measurements for each species and simulation on how low replicate numbers may affect coefficient of variations see Supplementary Section 1.

To compare intra- versus inter-specific variation we used coefficients of variation and Trait Probability Density (TPD) analyses (Carmona *et al.*, 2016). Coefficient of variation (CV), i.e. the standard deviation divided by the mean, defines the variability (or dispersion) of a set of values and was calculated using the raster package in R (Hijmans and van Etten, 2012). As CV describes the relative variability of the data, this value can be compared across different types of measurement. We determined intraspecific variation by calculating the CVs of each species for each trait independently and then taking the mean of these values (termed hereafter Intraspecific CVs). Interspecific variation was calculated by taking the mean trait value for each species and then calculating the CV across species (termed hereafter Interspecific CV; Jung *et al*. 2010). If the Interspecific CV is larger than the Intraspecific CVs, then the assumptions are met requiring larger interspecific than intraspecific variation for mean trait statistics. For species with fewer than 13-15 replicates, the CV may not be as reliable (see Supplementary Section 1), highlighting the need for future studies to make more measurements. To further explore the intraspecific variation, we used the TPD package in R (Carmona *et al.*, 2016) to extract a 2-dimensional Trait Probability Density (i.e. both size and density traits together) for each species. Trait values were logged (natural) prior to constructing the TPD to normalise the distribution. We then used TPDs to examine the overlap in trait values between species, species dissimilarity and the Rao functional diversity measure (de Bello *et al.*, 2013; Carmona *et al.*, 2016). Dissimilarity consists of both differences in trait space and differences in trait value probabilities within the same trait space, and the TPD package allows the decomposition of these two elements by calculating the overlap in shared trait space between species as part of the dissimilarity measure. Trait value probabilities are calculated as part of the TPD function when applied to species, essentially, the trait space has a total value of 1 and, within this space, trait values that are more likely to occur when and individual of that species is measured is given a higher value. This allowed us to explore whether dissimilarities were primarily due to differences in species traits, or differences in trait value probabilities across their TPD (Carmona *et al.*, 2016). TPDs were also used in the calculation of the Rao functional diversity, as overlap methods used with Rao can more accurately characterise functional diversity (de Bello *et al.*, 2013). All analyses were conducted in R (version 3.5, R Core Team, https://www.r-project.org/).

To aid the interpretation of the TPDs and trait spaces, we subdivided fungal species into three fruit body types: pileate, resupinate and half-resupinate (i.e. largely resupinate, but occurring with a small cap; Nordén *et al.*, 2013; Ryvarden and Melo, 2014). Resupinate and half-resupinate consisted of both polypore and corticioid species. Pileate (or bracket) fruit bodies were only polyporous species and we did not have occurrences of several brackets attached together.

## Results

Collected trait data on fruit bodies showed considerable variation prior to analysis. The largest fruit body was a red-listed half-resupinate polypore *Trichamptum laricinum* species that was 10932.74 cm^3^, covering the bottom and sides of an entire log, while the smallest was a minute 0.003 cm^3^ common corticoid resupinate *Veluticeps abietina*. The large *T. laricinum* fruit body was much larger than the second biggest fruit body (*Trichaptum abietinum*, 3013.601 cm^3^) and so CV values were calculated both with and without this large value. The half-resupinate perennial polypore *Phellinus viticola* had the densest recorded fruit body, with a dry density of 2.435 mg per mm^3^. The least dense fruit body was from the annual resupinate corticoid *Asterodon ferrugineofuscus* species and its dry density was only 0.006 mg per mm^3^.

CV values were enormous for fruit body size and large for fruit body density (Table 1). The Intraspecific CVs for fruit body size was smaller than the Interspecific CV, but not by much as their mean value was 77% of the Interspecific CV (Table 1). Fruit body density Intraspecific CVs value was more than double that of the Interspecific CV value (Table 1). This means that intraspecific variation was over two times more variable than interspecific variation in fruit body density.

**Table 1:** Coefficient of variation (CV) expressed as a percentage, for each species individually, the mean of species CVs and the CV of the community where a mean trait value was used for each species. The values in grey brackets below *T. laricinum* and the in the group CVs show the CV if the value for the exceptionally large *T. laricinum* fruit body was removed from the dataset. Shortened species names used in the figures below are also included in the table.

| Species or CV source | CV for fruit body size | CV for fruit body dry density |
| --- | --- | --- |
| *Antrodia serialis* (A. ser.) | 385.30 | 53.95 |
| *Asterodon ferruginosus* (A. fer.) | 146.61 | 146.58 |
| *Fomitopsis pinicola* (F. pin.) | 69.13 | 8.44 |
| *Fomitopsis rosea* (F. ros.) | 156.16 | 18.49 |
| *Gloeophyllum sepiarium* (G. sep.) | 156.94 | 39.57 |
| *Phellinus ferrugineofuscus* (P. fer.) | 261.70 | 26.38 |
| *Phellinus viticola* (P. vit.) | 122.98 | 152.35 |
| *Stereum sanguinolentum* (S. san.) | 236.74 | 64.80 |
| *Trichaptum abietinum* (T. abi.) | 206.61 | 51.87 |
| *Trichaptum fuscoviolaceum* (T. fus.) | 138.29 | 59.46 |
| *Trichaptum laricinum* (T. lar.) | 223.05<br>(197.43) | 48.16 |
| *Veluticeps abietina* (V. abi.) | 167.66 | 49.36 |
| Intraspecific CVs<br>(mean of above CVs) | 189.26<br>(187.13) | 59.95 |
| Interspecific CV<br>(CV of community using mean traits) | 244.90<br>(208.22) | 20.18 |

There was a large amount of overlap in the TPDs of species (Fig. 1), indicating that all species occupy a very similar trait space in terms of fruit body size and density. This was also demonstrated by the large amount of shared trait space calculated between species, with 24 out of the 66 species pairing combinations having a greater than 90% overlap and 11 species pairs having 100% trait overlap (Fig. 2). Despite this large overlap, there were dissimilarities greater than 0.5 for many species pairs (Fig. 2), due to the varying trait value probabilities of each species within the overlapping trait space (Fig. S2). This means that if both dissimilarity and trait overlap of a species pair was large, either the distribution of values within the shared space was different or the trait space of one species was larger than, and covering most of, the trait space of the second species (e.g. the latter case for *A. ferruginosus* and *F. pinicoloa*; Figs. 1, 2, S2). Pileate and resupinate fruit body types tended to have smaller trait space coverage than half-resupinates (with the exception of *A. ferruginosus*; Fig. 1B-D). The Rao functional diversity metric based on the TPDs was only 2.53, indicating that there is low functional distinctiveness between species.

**Figure 1.**
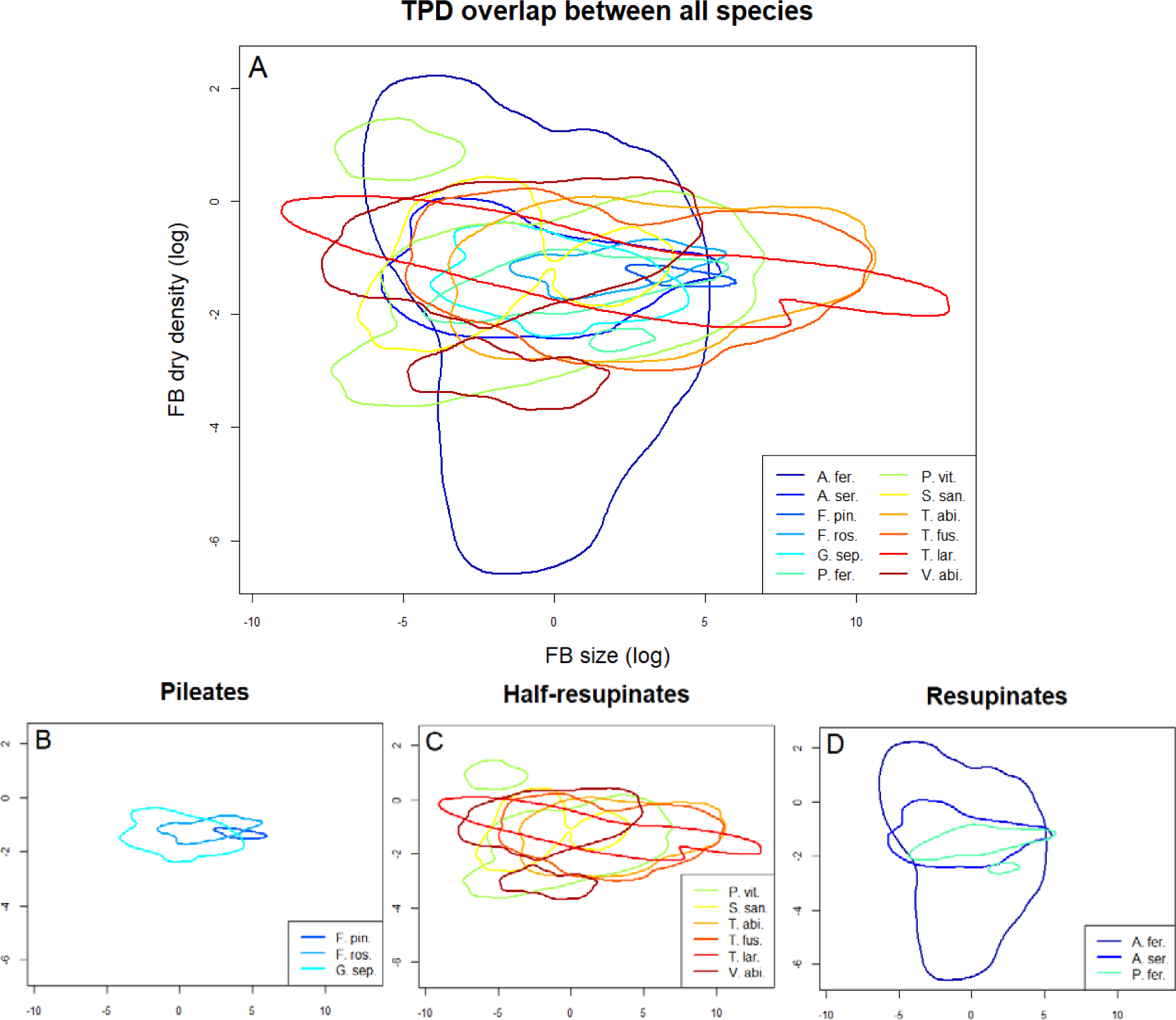
Trait Probability Density (TPD) outlines for the twelve fungal species; (A) demonstrates the degree over overlap across all species; the other figures are based on (A) but show only the species of a specific fruit body type, to enable easier viewing: (B) pileate fruit bodies, (C) half-resupinate fruit bodies and (D) resupinate fruit bodies. Note: the axis labels for (B)-(D) are the same as (A). The full TPD is shown for each species individually in Fig. S2. Full species names are shown in Table 1.

**Figure 2:**
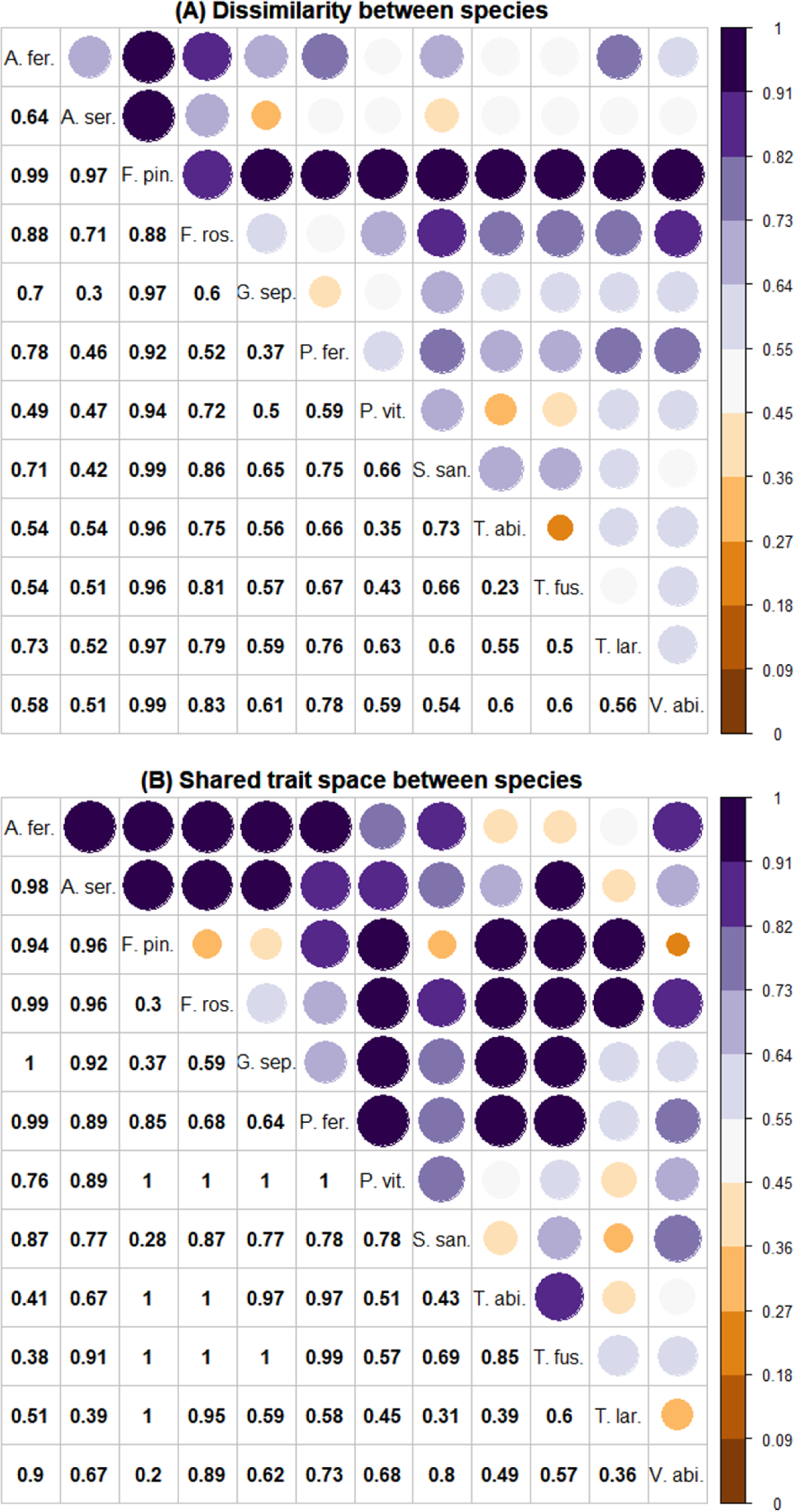
(A) Dissimilarity between species and (B) the proportion of shared trait space between species. Although species occupy much of the same trait space, differences in trait value probability across TPDs also contributes to the dissimilarity between species. Full species names are shown in Table 1.

Some individual species were notable for their small or extremely high variation. The common pileate polypore, *F. pinicola*, for example, had low variation overall (Table 1) and occupied the smallest trait space, but did have high probabilities of trait values occurring across this space (Fig. 1, Fig. S2). This meant that *F. pinicola* was the most dissimilar species in the analysis. *A. ferruginosus* and *P. viticola* both had highly variable fruit body densities. *A. serialis* had the highest CV for fruit body size (Table 1), *T. laricinum* covered the largest trait space on the x-axis (Fig. 1), the difference between the two being the single very large *T. laricinum* fruit body referred to previously.

## Discussion

Of the two fungal fruit body traits measured, density had more than a two-fold greater intraspecific variation than interspecific variation and size had enormous variation overall (on average 190 % dispersion around the mean), despite having slightly lower intraspecific variation. There was also a large amount of overlap in the trait space occupied by species and low functional diversity (Rao metric = 2.5) in relation to the number of species. Pileate fruit bodies tended to have the lowest variability, while half-resupinates were more variable. However, we have a relatively low number of measurements and there is an urgent need for more data to better characterise the trait space of fungal fruit bodies. Even where intraspecific variation was lower than interspecific, it was still exceptionally high, which has implications for how we interpret studies using species mean trait values. This is the first study to attempt to quantify the intra- to inter-specific variability in traits of wood fungi.

The physical size and density of fruit bodies clearly have important implications for the reproduction and dispersal of wood fungi (Aguilar-Trigueros *et al.*, 2015; Bässler *et al.*, 2015; Abrego *et al.*, 2017; Dawson *et al.*, 2018), but our measures may also depend on differences in the length of time that fruit bodies remains sufficiently intact. Pileates had the most restricted trait space, but there was a large amount of overlap between all species. Potentially pileates occupy smaller trait spaces because they are hardy and long-lived structures (with more skeletal hyphae in the case of *Fomitopsis* species). Whereas thinner and more fragile fruit bodies of half-resupinates and resupinates, like *A. serialis, P. ferrugineofuscus* and S. *sanguinolentum*, may vary more due to the shorter longevity of their fruit bodies (Jönsson *et al.*, 2008). These more short-lived species are likely more dependent on fast and effective resource allocation and fruiting when conditions are favourable (R-selected traits), whilst traits for coping with competition and stress may be more important than plasticity in fruiting in the more long-lived species (C- & S-selected; Boddy & Hiscox 2016). Hence, a greater variability in size and density among these short-lived half-resupinate and resupinate species may more tightly reflect responses in relation to environmental fluctuations or microsite environmental conditions. However, such relationships between trait variation and environmental conditions remain to be explored. One component that potentially could have accounted for the trait variation we saw was the volume of wood, given that it is the local food resource and important for species composition (Edman *et al.*, 2004; Nordén *et al.*, 2013). However, in this case, wood volume either had no discernible relationship (fruit body density) or a statistically significant, but very low explanatory power relationship (R^2^ of 0.03; Fig. S3), meaning that volume did not explain much of the variation in fruit body traits. Therefore it is unlikely that the trait variations we observed were due to wood volume.

We showed that the variation in fungal fruit bodies is overall very high, with a large amount of overlap in trait space. Fruit body size in particular had an extremely large amount of variation compared with other organisms, where CV values were in the 10s (or less) rather than 100s (e.g. fish: Meiri *et al*. 2005; carnivore skull morphology: Blanck & Lamouroux 2006; bird sperm length: Kleven *et al*. 2008), or plants, where CVs are generally below 10 (e.g. Jung *et al*. 2010; Messier *et al*. 2017). When comparing with mycorrhizal studies, CV values of fruit body density were similar to fungal biomass (Johnson *et al.*, 2012), but in the mycorrhizal case, intra- and inter-specific variation were roughly equivalent. The fungi studied here have indeterminate growth size, meaning that they may be more variable. It is possible that fruit bodies of other types of fungi (e.g. Agaricomycete soil fungi), may have less variation than the wood fungi we studied, however a recent examination of fungal taxonomic books from different countries showed considerable variation in saprotrophic and ectomycorrhizal agaric species (Halbwachs & Karasch, 2019). Within species variation in this study had cap diameter CV values of up to 35 which, given these are the values used to identify species, indicates that agaric or mushroom forming species are also likely to have considerable intraspecific variation. Data on variation in these species is needed to test this (Halbwachs & Karasch, 2019). While we did not discriminate fruit bodies based on the ‘age’ of the fruit body, as this is difficult to determine, the largest two fruit bodies were from annual species (Nordén *et al.*, 2013). Despite this restriction, the very high degree of variation is undeniable, which has implications for future research. If mean trait values are to be used (as they undoubtedly will), they should be measured on site, rather than sourced elsewhere. This will help ensure they are reflecting local processes more accurately.

Another concern is the high overlap of trait space among species, supported by the low Rao value, and the implications this has for understanding species niches. Species with a low degree of overlap or high functional distinctiveness occupy different functional niches and are likely to be performing different roles in the ecological community, whereas the opposite is true when there is high functional redundancy (i.e. trait space overlap and low Rao value; Carmona *et al*. 2016). The strong functional redundancy in our data indicates that if niche partitioning is operating, it may be at the individual fungus and log level, rather than the species. A study examining intraspecific variation in the ectomycorrhizal fungus *Laccaria bicolor* found there was niche partitioning at the strain (individual) level where nutrients were limited (Hazard *et al.*, 2017). Niche theory predicts that filters determine trait (and classically therefore species) composition at plot level, however a central tenant of the opposing neutral theory is that species identity is irrelevant (Messier, McGill and Lechowicz, 2010). Therefore fungi present an interesting example in the debate between these two theories, if niche partitioning at the individual level is occurring. It is vital that more individuals are measured at the site level to determine if fungi, though highly plastic, do partition themselves into separate trait space at the log or plot level (i.e. using methods such as; Ackerly & Cornwell 2007). We were unable to determine this from our data as we only took measurements from one log per plot and only three fruit bodies per species on that log.

Intraspecific variation was not only high in our study, but for fruit body density it was more than double interspecific variation. This high within species variability has implications for how we interpret and use trait values in future fungal studies (Violle *et al.*, 2012; Des Roches *et al.*, 2018). Before we highlight these issues, we should emphasise that more data are needed on all fungal traits to determine if these trends hold or are unique to fungal fruit bodies, particular to wood-decay fruit bodies, or even just these two traits. Although the fruit body traits chosen here may be more variable then other traits, such as spore size (thought to be more conservative), there are indications that even these traits may still have high variability. Halbwachs & Karasch (2019) show that intraspecific variation for spore traits just between identification book values (i.e. the taxonomic guides of different countries) can have CVs as high as 42, indicating that the true intraspecific variation will be much higher. While mean trait value studies are useful for gaining a more general understanding, results will need to be interpreted with caution if the high variability is not taken into account (Jung *et al.*, 2010; Violle *et al.*, 2012; Shipley *et al.*, 2016; Des Roches *et al.*, 2018). Special care is needed if mean trait values are used to identify underlying ecological mechanisms, characterise species niches or persistence in a community (Violle *et al.*, 2012). Even in plant species, when variation was low and intraspecific variation was at least 2.8 times smaller than interspecific variation, Jung *et al*. (2010) found that including intraspecific variation could account for up to 44% of the trait-environment gradient. Given the variation indicated by our data, it would be interesting to examine in the future, how much intraspecific variation is contributing to trait-environment interactions. However, it is possible that the very high intra- to inter-specific in our case may be inflated by two potential causes: first, as mentioned previously, we were unable to determine fruit body ‘age’ which may affect density; and secondly we only have 12 species in our dataset, some with few measurement values, so further data collection from other species may increase or decrease the variation. This is especially the case in species with low measurement numbers, such as *A. ferruginosus* or *T. laricinum*, where the CV values will likely change as more data is collected. Even if both these causes had some effect in this case, there is still a strong indication that intraspecific variation is very high within fungal fruit body traits. Although we did sample multiple fruit bodies from the same log, if these were the same individual it would lead to underestimation of intraspecific variation, which is the opposite to the issue we highlight here. Further data collection is vital to future fungal trait studies examining fruit body traits, and these projects should aim to have at least 13-15 measurements per species, although this recommendation may change as the characteristics of variation become more known.

Shipely *et al*. (2016) point out that if species are so variable that individual trait measurements are needed then trait ecology loses the asset of being generalizable beyond the species. However, in the case of fungi, if traits are as variable as is widely believed, there are two reasons to continue collecting individual traits. First, the ecology of most fungi is still poorly known and through measuring traits at the individual level we will gain a deeper understanding of species and fungal ecology (i.e. not just the Raunkiæran Shortfall, but also the Hutchinsonian and Eltonian Shortfalls; Hortal *et al*. 2015). Secondly, if fungal species have large trait space overlap, as is indicated in this study, then species identity may not be as much use in understanding niche space and community dynamics as in other organisms, such as plants. Therefore, by measuring traits and their variability at fine (i.e. the wood unit) scales we may be able untangle community dynamics in a more meaningful way than if species identity alone is used.

Our study has demonstrated both high overall trait variation and similarly high intraspecific variation in wood-decay fruit body traits. Trait ecology is a burgeoning field in which fungal ecologists are only just beginning to participate (Crowther *et al.*, 2014; Aguilar-Trigueros *et al.*, 2015; Dawson *et al.*, 2018). However, by taking theory and techniques made on assumptions that are generally true for plants, we may be missing important processes in highly variable organisms like fungi. It is essential that future studies measure traits so that there is a sound quantitative understanding of both intra- and inter-specific variation. We found that intraspecific variation, in the case of fruit-body density, was more than twofold interspecific variation, and even in fruit-body size, differences between the two was negligible compared with overall variation. Further trait space overlap was also extremely high, with species dissimilarities largely due to differences in species trait value probability distribution.

## Acknowledgements

We would like to thank Carlos Perez Carmona for his assistance with the TPD package and metrics. Also we would like to thank the field assistants: Elisabet Ottosson, Håkan Berglund, Torbjörn Josefsson and Sofia Nygårds. Finally we would like to thank the two anonymous reviewers, two others who gave feedback on the preprint: Franz-Sebastian Krah and Hans Halbwachs, and Lynne Boddy for editing. The study was funded by the Swedish Agricultural University to S.K.D. and FORMAS Grant 2016-00461 to S.K.D. and M.J.

## Supplementary Section 1: Measurement numbers and effect on coefficient of variation

In our article we present the coefficient of variation (CV) values for 12 different saprotrophic fungi species. However, for some species we have a low number of fruit body measurements (Table S1). For these species, especially *A. ferruginosus* or *T. laricinum*, further data may change the CV values presented to be somewhat lower or higher.

**Table S1:** the number of fruit body measurments for each species presented in the article.

| Species or CV source | Number of measurements |
| --- | --- |
| *Antrodia serialis* (A. ser.) | 25 |
| *Asterodon ferruginosus* (A. fer.) | 5 |
| *Fomitopsis pinicola* (F. pin.) | 7 |
| *Fomitopsis rosea* (F. ros.) | 11 |
| *Gloeophyllum sepiarium* (G. sep.) | 17 |
| *Phellinus ferrugineofuscus* (P. fer.) | 17 |
| *Phellinus viticola* (P. vit.) | 15 |
| *Stereum sanguinolentum* (S. san.) | 6 |
| *Trichaptum abietinum* (T. abi.) | 13 |
| *Trichaptum fuscoviolaceum* (T. fus.) | 7 |
| *Trichaptum laricinum* (T. lar.) | 5 |
| *Veluticeps abietina* (V. abi.) | 18 |

To demonstrate the effect of the number of measurements on CV we can examine how CV values change with the number of ‘measurements’ drawn from a randomly generated dataset. In Fig. A1 we present a randomly generated dataset of 1000 ‘measurements’ with a mean of 5 and standard deviation 1. We then show the variation if we take a draw of 5 ‘measurements’ from this dataset 100 times. Finally, we show how the variation in CV changes as you increase the number of measurements used. This shows that for a randomly generated dataset, variation in CV stabilises somewhat after 13-15 measurements and then again after around 50. In regards to this paper, it shows that species with low numbers of measurements may change in CV values as more data is recorded. For fungi more generally, if variation follows the same structure as this randomly-generated dataset, then at least 13-15 measurements are needed, however this number may change as more information is collected and the structure of fungal variation known.

**Figure S1:**
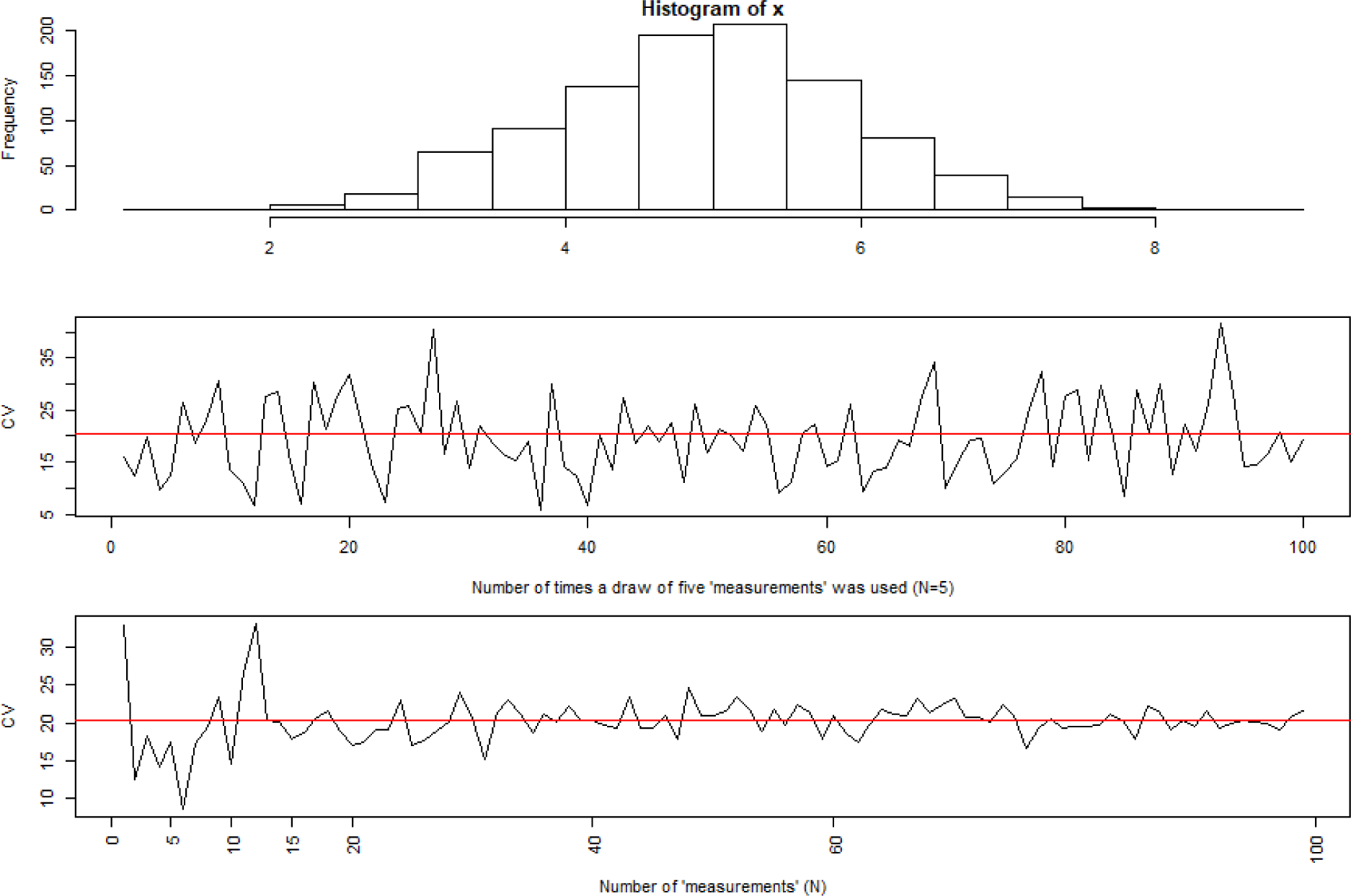
Plots showing the randomly generated dataset and the effect of the number of measurements taken on CV; red lines indicate the CV of the whole dataset. R code for generating these plots is on the following page. ~~~
#code to simulate the effect of number of replicates on measurement of coefficients of variation
(CV)

library(raster)

#create a dataset of random values and then the data for each plot below
x <- rnorm(n = 1000, mean = 5, sd = 1)

#set up plotting
par(mfrow = c(3,1), mar = c(4,4,1,0))

#distribution of x
hist(x, xlab = “”)

#plot showing the variation in CV when only 5 values from X are used (equivalent to calculating CV
from 5 measurements of fruit bodies), red line shows CV value for whole dataset

plot(sapply(rep(5, 100), function(i) cv(sample(x, i))), type =“l”,
   xlab=“Number of times a draw of five ‘measurements’ was used (N=5)”, ylab=“CV”)
abline(h = cv(x), col = ‘red’)

#plot showing how the variation of CV changes as you increase the number of ‘measurements’
xticklabels = c(0, 5, 10, 15, 20, 40, 60, 100)
plot(sapply(2:100, function(i) cv(sample(x, i))), type =“l”,
   xlab=“Number of ‘measurements’ (N)”, ylab=“CV”, xaxt=‘n’)
axis(1, at = xticklabels,las=2)
abline(h = cv(x), col = ‘red’)
grid()
~~~

**Figure S2:**
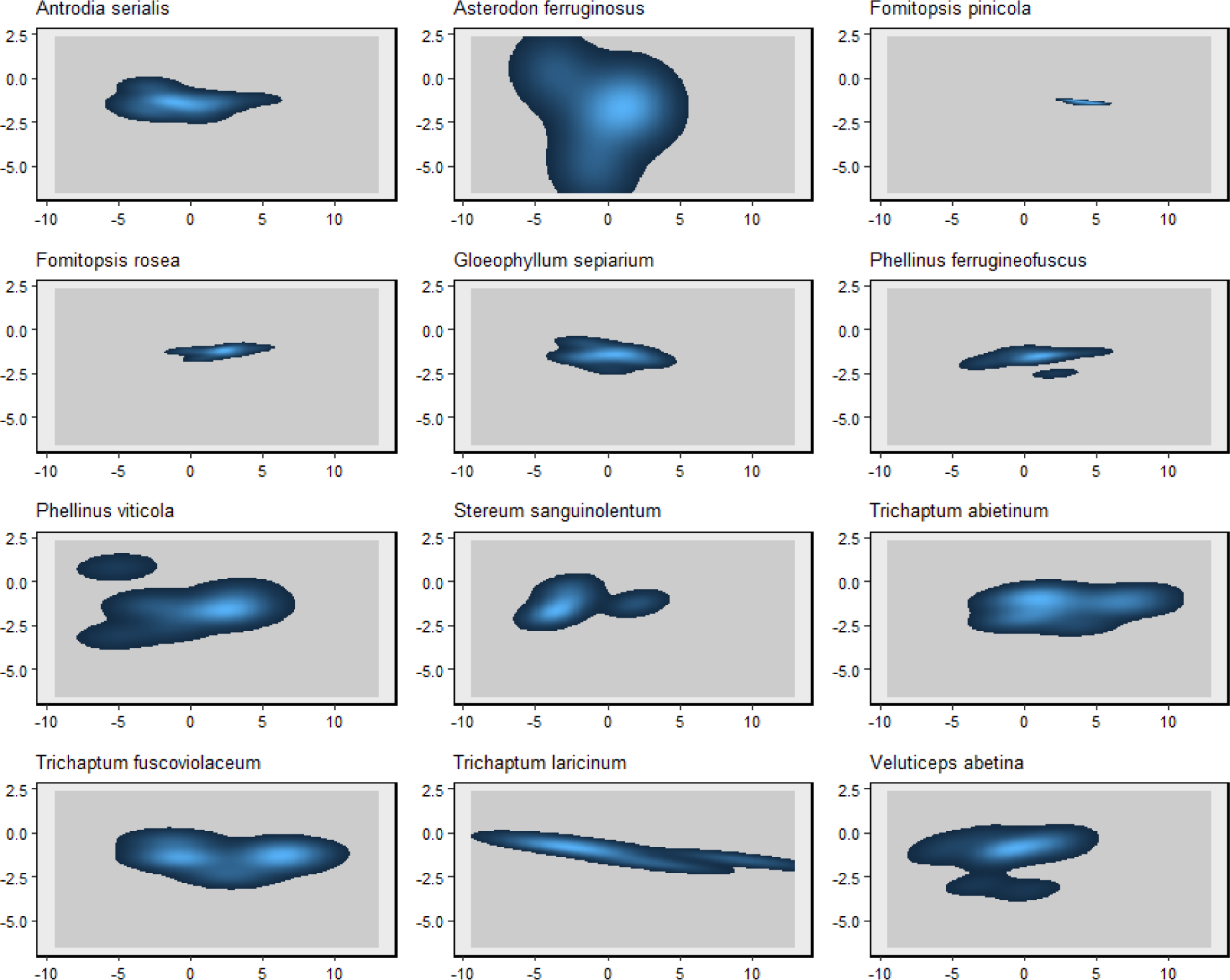
Trait probability densities for the twelve fungal species with abundance distribution shown. Full species names are shown in Table 1. Lighter blue indicates a higher probability of that combination of values occurring when that species is measured, with each TPD space totalling one.

**Figure S3:**
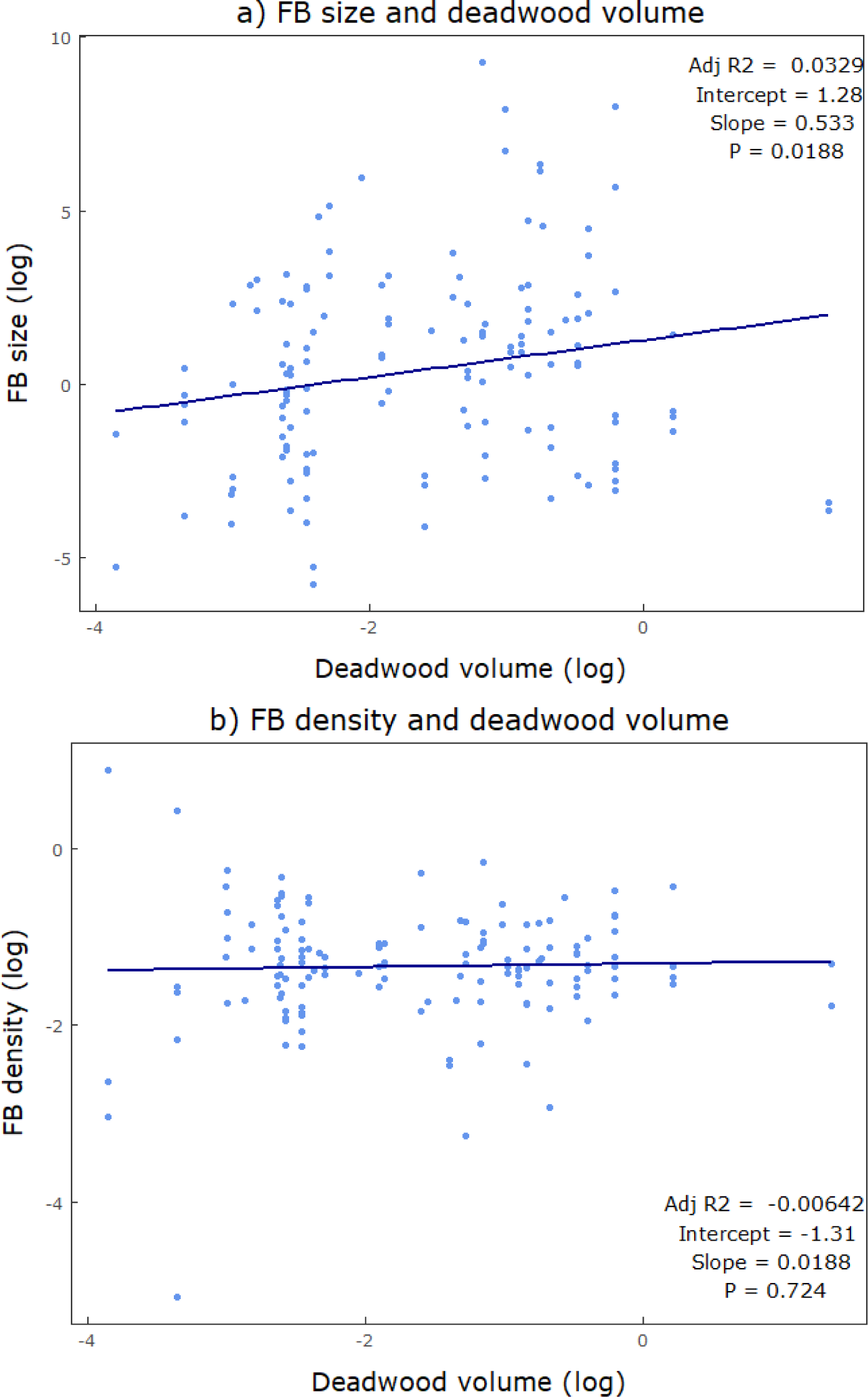
The relationship between each trait and the volume (m^3^) of the wood resource that trait was measured on. All values were log transformed to normalise distribution. There was no significant relationship between fruit body density and wood volume, but fruit body size had a significant, though very low explanatory power relationship.

